# A comparison of shear- and compression-induced mechanotransduction in SW1353 chondrocytes

**DOI:** 10.1101/2021.05.25.445657

**Authors:** Hope D. Welhaven, Carley N. McCutchen, Ronald K. June

## Abstract

Mechanotransduction is a biological phenomenon where mechanical stimuli are converted to biochemical responses. A model system for studying mechanotransduction are the chondrocytes of articular cartilage. Breakdown of this tissue results in decreased mobility, increased pain, and reduced quality of life. Either disuse or overloading can disrupt cartilage homeostasis, but physiological cyclical loading promotes cartilage homeostasis. To model this, we exposed SW1353 cells to cyclical mechanical stimuli, shear and compression, for different durations of time (15 and 30 min). By utilizing liquid chromatography-mass spectroscopy (LC-MS), metabolomic profiles were generated detailing metabolite features and biological pathways that are altered in response to mechanical stimulation. In total, 1,457 metabolite features were detected. Statistical analyses identified several pathways of interest. Taken together, differences between experimental groups were associated with inflammatory pathways, lipid metabolism, beta-oxidation, central energy metabolism, and amino acid production. These findings expand our understanding of chondrocyte mechanotransduction under varying loading conditions and time periods.

## Introduction

Mechanotransduction is the study of the cellular responses to applied mechanical loads and deformations (Paluch et al., 2015). In the past, mechanotransduction has been studied in sensory cells, such as hair cells showing intricate mechanisms of mechanosensing (Gillespie and Muller, 2009). Beyond sensory cells, the musculoskeletal system, consisting of bone, cartilage, muscle, tendons, and ligaments, functions to provide support, stability and movement for the organism. In doing so, these tissues experience a variety of mechanical loads (compressive, tense, and shear) that, through cellular mechanotransduction, must be interpreted, dispersed, and transduced by resident cells. Physiological levels of loading are necessary for cell growth, proliferation, and survival in tissues like articular cartilage (AC) (Jaalouk and Lammerding, 2009; Orr et al., 2006).

AC provides near-frictionless articulation on the ends of long bones which provides a smooth surface for joint movement, resists extreme loads that are applied to the joint, and reduces the stress placed on underlying bone (Sophia Fox et al., 2009). Chondrocytes synthesize an extracellular matrix (ECM) which is the main component of cartilage (Sanchez-Adams et al., 2014). The ECM performs mechanotransduction and helps to maintain homeostasis during mechanical loads. Therefore, under load associated with joint motion, the ECM is stimulated to support cartilage homeostasis. Although sub-injurious mechanical loading is required to maintain cellular homeostasis, loads large in duration and/or magnitude can lead to biological imbalances that disrupt homeostasis and induce joint disease such as osteoarthritis (OA) (Racunica et al., 2007).

OA is a chronic joint disease that is caused by many factors including inflammation and joint injury that results in the eventual breakdown of AC. Degradation of tissue ultimately leads to joint pain, limited mobility, and reduced quality of life. Therefore, investigation of mechanotransduction and metabolism of AC, chondrocytes, and the ECM may elucidate the effects of mechanical loading on the joint.

To this end, there has been substantial research in the field to expand our understanding of how chondrocytes respond to the mechanical stimuli experienced by the joint (Bricca et al., 2017; Chan et al., 2018; Chan et al., 2016). Building on these results, this study examined how chondrocytes respond to cyclical shear and compressive loading *in* vitro by utilizing untargeted metabolomics.

Metabolomics is an analytical profiling technique used to investigate large numbers of small molecule intermediates called metabolites. These molecules “act as a spoken language, broadcasting signals from the genetic architecture and the environment (Jewett et al., 2006).” Metabolomic profiling analyzes thousands of small molecules characterizing the cellular phenotype and provides an unbiased view of metabolic shifts induced by experimental conditions (Carlson et al., 2018). This technique is applicable to the study of chondrocyte metabolism and OA because it has the ability to provide metabolic information, such as involved pathways and intermediates, in response to applied stimuli, such as shear and compression. While many studies have investigated the effects of mechanical stimuli on the joint, few have investigated if mechanical stimuli impact the metabolism of involved tissues.

Several studies focused on the health of cartilage, changes in the microenvironment, and structure of the ECM after exposure to various levels of mechanical stimuli (Bricca et al., 2017; Chan et al., 2018; Chan et al., 2016). Brica *et al* varied doses of exercise and revealed that cyclical loading serves as a protective tool for chondrocytes to withstand loads that are increasing in magnitude and frequency (Bricca et al., 2017). In contrast, extended periods of increasing load can interfere with homeostasis leading to deterioration of the ECM which can induce joint disease such as OA (Chan et al., 2018). Additional studies measured mechanical strain in cartilage and the effects of compressive loading with non-invasive imaging and found localized changes in the cartilage mechanical environment that ultimately led to cartilage deformations (Chan et al., 2018; Chan et al., 2016). To study the mechanobiology of these load-bearing tissues, varying loading frequency and amplitude were applied to bone and cartilage explants *in vitro* through a dual frequency system. This study revealed that cyclical compression during these activities stimulates cellular signaling and chondrocyte metabolism by tracking solute transport (Chan et al., 2016).

Although progress has been made in understanding chondrocyte mechanotransduction, many questions remain. Additional data is needed to understand how chondrocytes respond to shear and compressive loading. Therefore, the objective of this study is to identify mechanosensitive differences between shear and compressive stimulation after 0, 15, and 30 minutes of 5 +/− 1.9% cyclical strain at 1.1Hz for SW1353 chondrocytes, encapsulated in physiologically stiff agarose. After loading, metabolite extracts were analyzed by liquid-chromatography-mass spectrometry (LC-MS) in search of metabolites that differentiate between the type and duration of loading. These data provide insight into how SW1353 chondrocytes respond to mechanical stimuli thus expanding our knowledge of chondrocyte mechanotransduction.

## Materials and Methods

### SW 1353 chondrocyte culture and encapsulation

SW1353 chondrocytes were cultured in DMEM with 10% fetal bovine serum and antibiotics (10,000 I.U./mL penicillin and 10,000 μg/mL streptomycin). Cells were first expanded in monolayer for one passage at 5% CO_2_ at 37°C prior to gel encapsulation. For encapsulation, cells were seeded in physiologically stiff agarose (4.5% v/v, Sigma: Type VII-A A0701) at a concentration of 500,000 cells/hydrogel (diameter: 7mm, height: 12.7mm). Hydrogel constructs were individually placed in wells, submerged in DMEM with 10% fetal bovine serum, and allowed to equilibrate for 24 hours prior to mechanical stimulation.

### Mechanical Stimulation

Cell encapsulated hydrogels were randomly assigned to five experimental groups (n=5 per experimental group): unloaded controls (0 minutes of mechanical stimuli), 15 or 30 minutes of cyclical shear, or 15 or 30 minutes of cyclical compression. Hydrogels were placed in DMEM with 10% fetal bovine serum without antibiotics and mechanically stimulated with cyclical shear or compression using a custom-built sinusoidal loading apparatus. Sinusoidal compressive and shear strains of 5.00±1.90% (based on initial 12.70mm ± 0.01mm gel height and 7.00mm ± 0.01mm gel diameter, respectively) were applied at 1.1 Hz to simulate the preferred stride rate of humans for 0, 15 and 30 minutes (Jutila et al., 2014; Jutila et al., 2015). All hydrogel mechanical testing was performed in physiological cell culture conditions (5% CO_2_, 37°C).

### Metabolite Extractions

Immediately after mechanical stimulation, hydrogels were flash frozen in liquid nitrogen, pulverized and placed in 70:30 (v/v) methanol : acetone. Samples were vortexed for 5 minutes and placed in −20°C for 5 minutes. This two-step process was repeated 4 times and samples were stored overnight (−20°C) for macromolecule precipitation. The next day proteins and other macromolecules were pelleted by centrifugation. The supernatant containing the metabolites was transferred to a separate tube and dried by vacuum concentration for 6.5 hours to remove solvents. Dried metabolites were resuspended in 100μL mass spectrometry grade 50:50 (v/v) water : acetonitrile solution immediately prior to high performance liquid chromatography-mass spectrometry (HPLC-MS) analysis.

### Untargeted Metabolomic Analysis

Metabolomics is the analysis of small molecules (metabolites) in a biological system that provides a global description of cellular function at a given point in time. Extracted metabolites were analyzed using HPLC-MS (Agilient 6538 Q-TOF mass spectrometer) in positive mode (resolution: ~20ppm, accuracy: ~5ppm, possible ionization adducts: H^+^, Na^+^). Peak intensities for m/z values in the experimental sample set were identified and exported using Agilient MassHunter Qualitative Analysis software. All data was log transformed and autoscaled prior to analysis. All statistical analyses utilized the transformed and scaled data using MetaboAnalyst (Xia and Wishart, 2016) (See Supplementary Materials and Methods).

## Results

A total of 1,457 distinct metabolite features were detected across all experimental groups. First, all experimental groups (control, 15 min. compression and shear, 30 min. compression and shear) were analyzed (Fig. 1). HCA and PCA examined overall variation between groups in the dataset. Neither shows complete separation between experimental groups (Fig. 1A-B), although the first two principal components are associated with more than 30% of the overall variance indicating moderate underlying structure in the dataset. To further examine this dataset, we applied PLS-DA to assess differences between experimental groups. Separation between experimental groups was observed with minimal overlap (Fig. 1C) using this supervised method.

**Figure 1.**
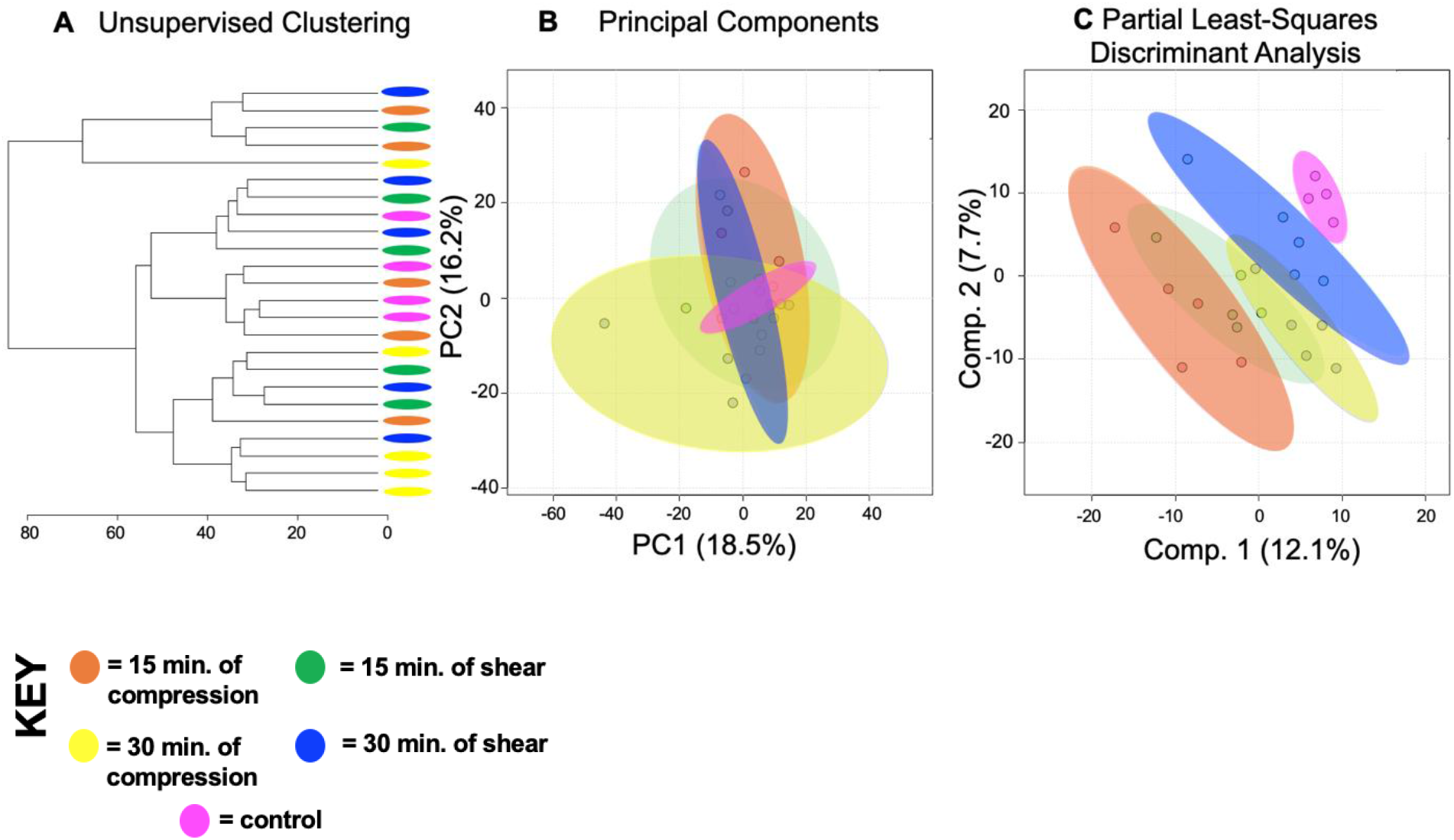
SW1353 chondrocytes have shared and distinct responses to shear and compressive forces. A total of 1,457 metabolites were analyzed by both unsupervised hierarchical clustering (HCA) and principal component analysis (PCA) and supervised partial least-squares discriminant analysis (PLS-DA). (A). Unsupervised HCA visualized by a dendrogram did not distinguish distinct clusters between the five experimental groups (control, 15. Min compression and shear, 30 min. compression and shear). HCA assigns Euclidean distances to illustrate dissimilarity between samples and reveal clusters of samples with similar metabolomic profiles. (B) PCA, similar to HCA, did not display distinguished clusters between the five experimental groups. PCA is shown as a scatterplot with the first two PC on the x and y axes. The x axis shows PC1, which accounts for 18.3% of the variation in the dataset. PC2 is on the y-axis and accounts for 14.2% of the variation in the dataset. (C) Supervised PLS-DA finds some separation between the five experimental groups, with similarity between different time points and different forces. PLS-DA is shown as a scatterplot of the top two components, with component 1 accounting for 14.6% and component 2 accounting for 7.6% of the variation in the dataset. The colors in A-C correspond to sample cohorts: pink – control, orange – 15 minutes compression, yellow – 30 minutes compression, green – 15 minutes shear, blue –30 minutes shear.

To compare the effects of shear and compression on metabolomic profiles, samples exposed to each loading type, shear and compression, were analyzed separately (Figs. 2-3). When analyzing differences between compressive samples between 0, 15, and 30 minutes of loading, HCA and PCA find clustering of samples within their respective timepoints indicating chondrocyte mechanotransduction (Fig. 2A-B). PLS-DA shows further discrimination with all samples clustering within their respective cohorts (Fig. 2C). We found similar results for shear stimulation. HCA and PCA revealed clustering of samples within each time point (Fig. 3A-B). PLS-DA shows further separation between samples from different shear groups (Fig. 3C). These data show that SW1353 chondrocytes respond to both shear and compression with alterations in their metabolomic profiles.

**Figure 2.**
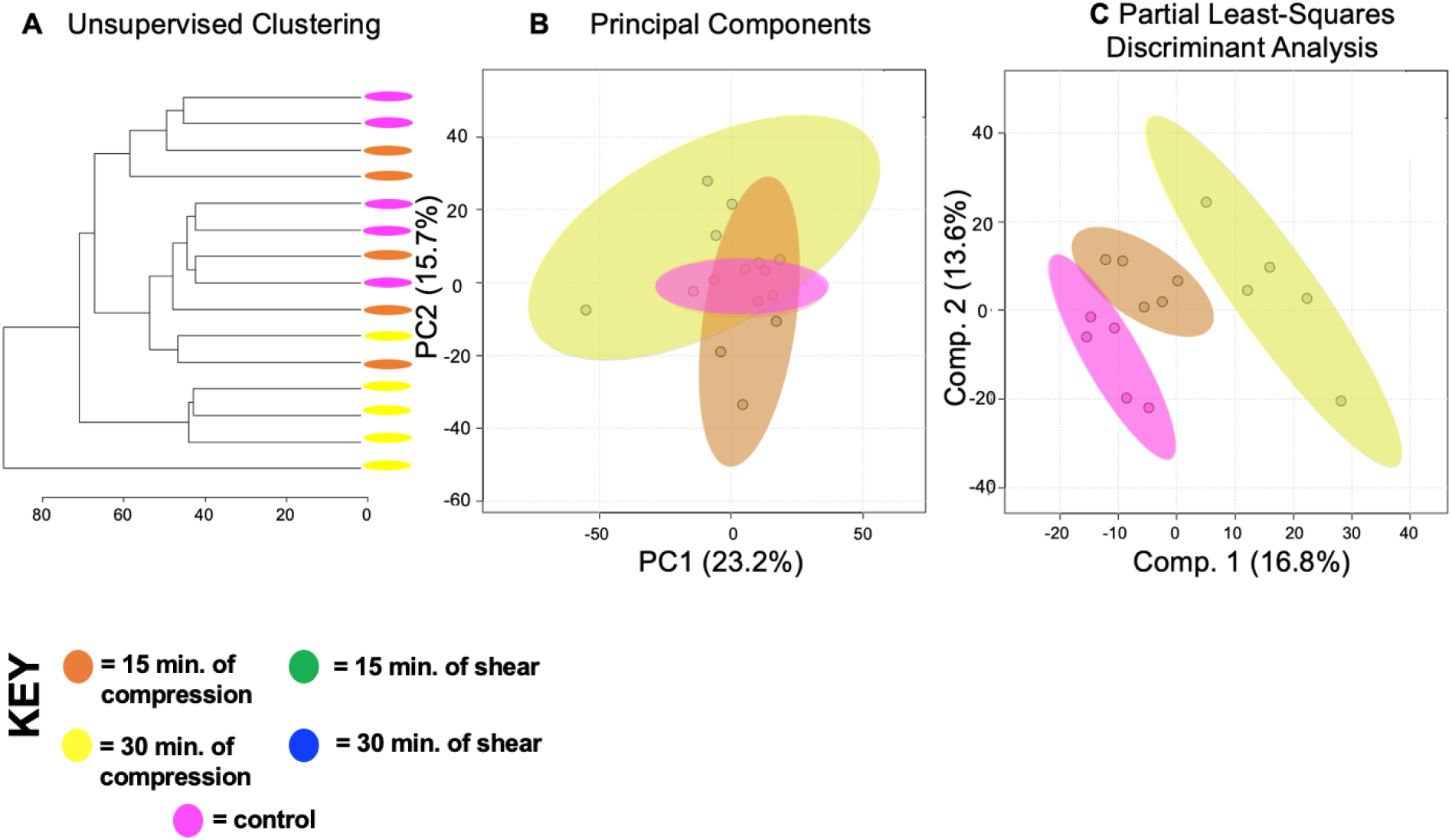
Differing amounts of exposure to compression reveal distinct metabolic profiles. Comparison of samples exposed to compression reveal metabolomic profiles of chondrocytes between 15 and 30 minutes of compression differ. (A) Samples compressed at different times separate into distinct clusters by HCA as illustrated in the dendrogram. (B) PCA displays some clustering of samples within their respective cohorts: chondrocytes exposed to compressive forces for 15 minutes (orange) and 30 minutes (yellow). Control chondrocytes that weren’t exposed to mechanical stimuli is displayed for comparison purposes (pink). PCA is shown as a scatterplot of the first two PCs (PC1 and PC2), which account for 23.2% and 15.7% of the overall variation in the dataset, respectively. (C) PLS-DA finds clear separation between samples. PLS-DA is visualized as a scatterplot of the top two components, which account for 16.8% and 13.6% of the overall variation in the dataset.

**Figure 3.**
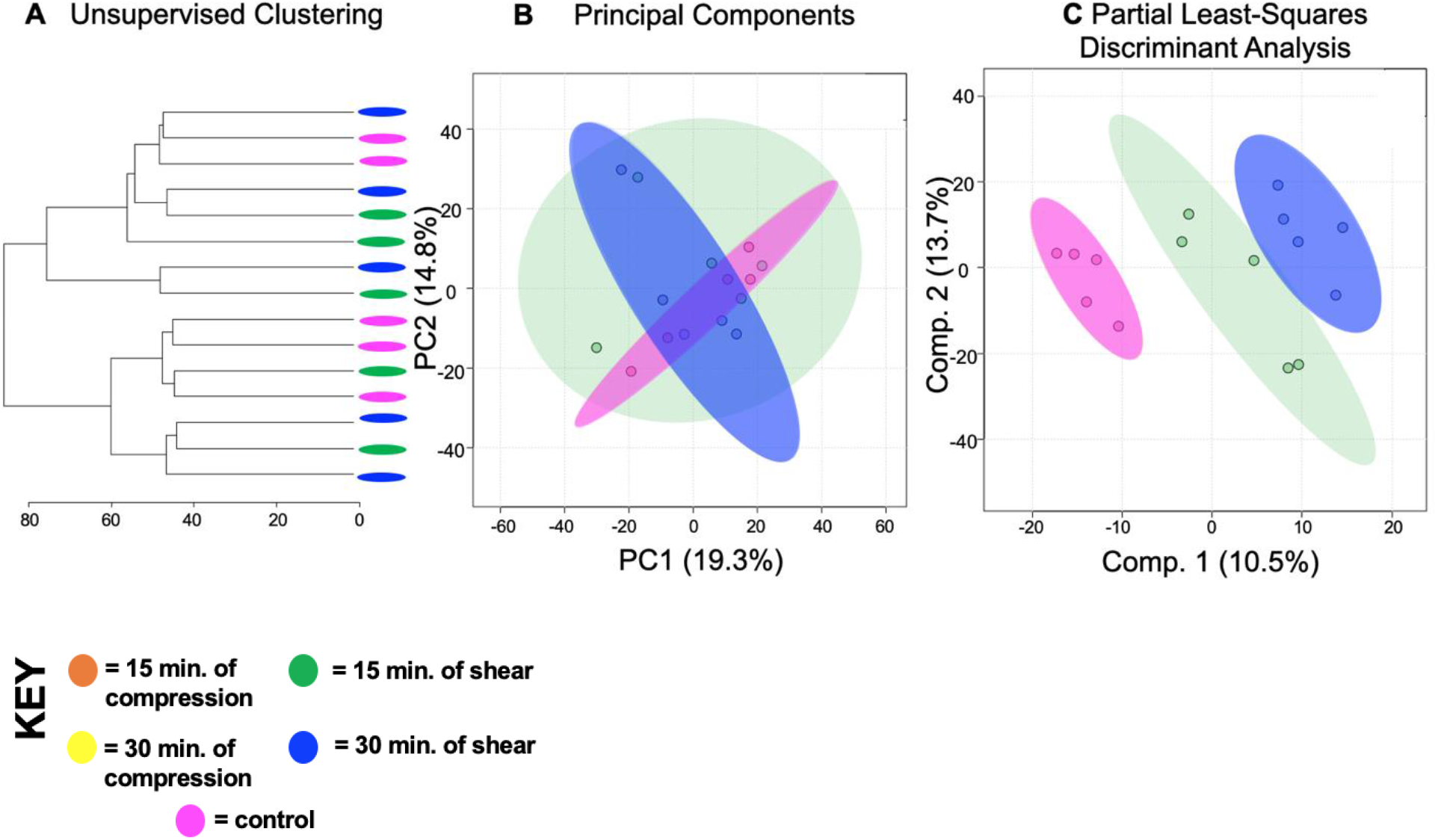
Differing amounts of shear force reveal distinct metabolic profiles. Metabolomic profiles of chondrocytes exposed to 15 and 30 minutes of shear force differ. (A) Samples compressed at different times separate into weakly distinct clusters by HCA as illustrated in the dendrogram. (B) PCA displays some clustering of samples within their respective cohorts: chondrocytes exposed to shear force for 15 minutes (green) and 30 minutes (blue). Control chondrocytes that weren’t exposed to mechanical stimuli is displayed for comparison purposes (pink). PCA is shown as a scatterplot of the first two PCs (PC1 and PC2), which account for 19.3% and 14.8% of the overall variation in the dataset, respectively. (C) PLS-DA finds clear separation between samples. PLS-DA is visualized as a scatterplot of the top two components, which account for 10.5% and 13.7% of the overall variation in the dataset.

Chondrocytes exposed to shear and compression for the same amount of time showed interesting similarities and differences between types of mechanical stimulation. When comparing compression and shear at 15 minutes, HCA and PCA showed limited separation and clustering (Fig. 4A-B). But PLS-DA clearly distinguishes samples between shear and compression at the 15 minute timepoint (Fig. 4C). Similarly, compression and shear samples that were both exposed to 30 minutes of stimulation were compared. Both HCA and PCA found clear separation and clustering of samples at 30 minutes (Fig. 5A-B) that was confirmed by PLS-DA (Fig. 5C). With this knowledge, we used VIP scores from PLS-DA and volcano plot analysis to identify specific metabolites that contributed to the differences between experimental groups. Specific metabolite features of interest were then matched to metabolite identities and pathways using MetaboAnalyst. Significant pathways consist of sialic acid metabolism (all experimental groups); prostaglandin formation from arachidonate, fatty acid metabolism, tyrosine metabolism (15 min. of compression vs 30 min. of compression); porphyrin metabolism (15 min. of shear vs 30 min. of shear); carnitine shuttle (15C vs. 15S); squalene and cholesterol biosynthesis (30C vs. 30S).

**Figure 4.**
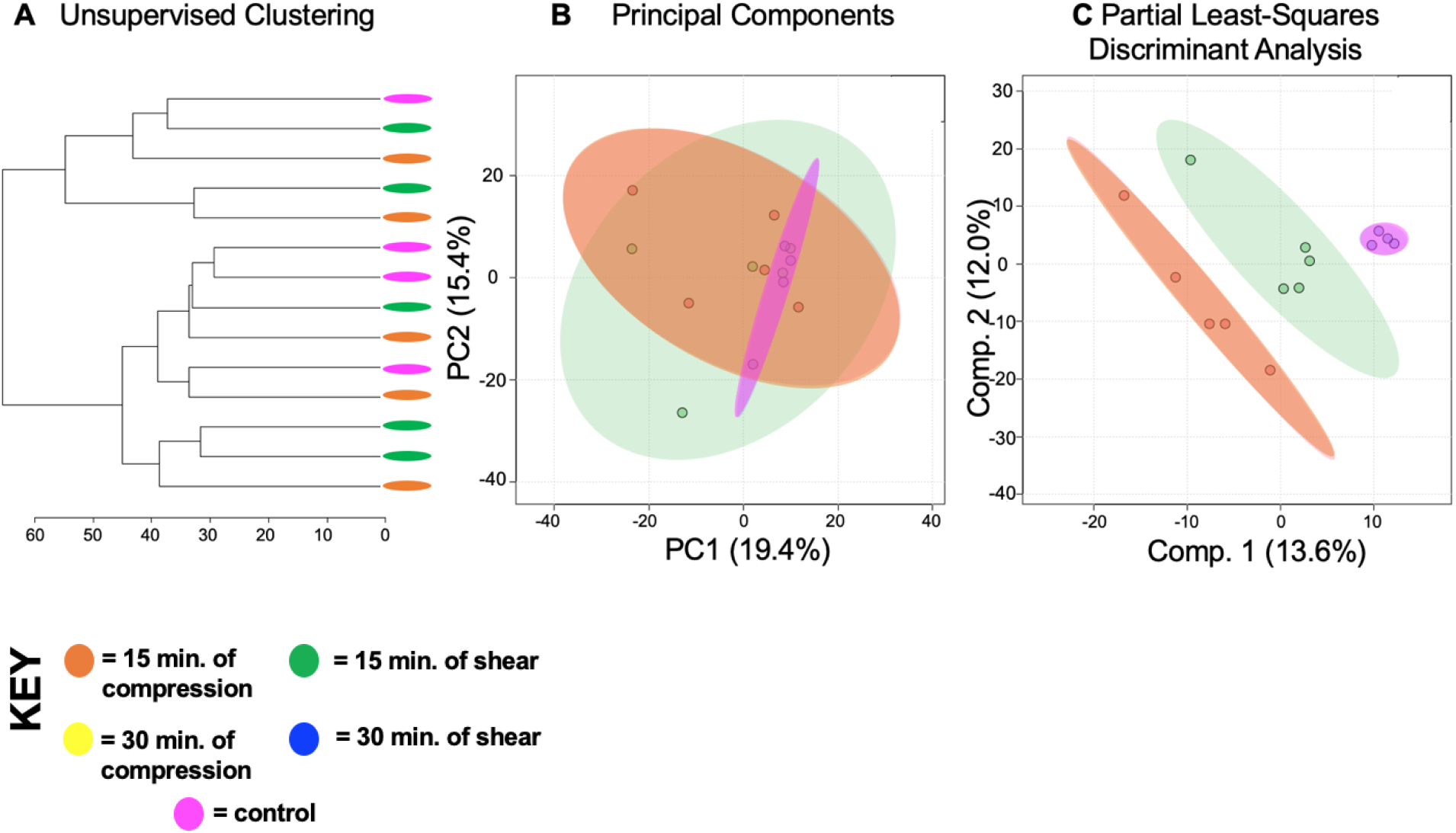
Metabolic profiles of differing forces with an equal exposure time of 15 minutes vary. Metabolomic profiles of chondrocytes exposed to compression and shear forces for 15 minutes differ metabolically(A). Samples exposed to different forces for the same amount of time somewhat separate into clusters by HCA as illustrated in the dendrogram. (B) PCA displays some clustering of samples within their respective cohorts: chondrocytes exposed to compression for 15 minutes (orange) and chondrocytes exposed to shear forces for 15 minutes (green). Control chondrocytes that weren’t exposed to mechanical stimuli is displayed for comparison purposes (pink). PCA is shown as a scatterplot of the first two PCs (PC1 and PC2), which account for 19.4% and 15.4% of the overall variation in the dataset respectively. (C) PLS-DA finds clear separation between samples. PLS-DA is visualized as a scatterplot of the top two components, which account for 13.6% and 12.0% of the overall variation in the dataset.

**Figure 5.**
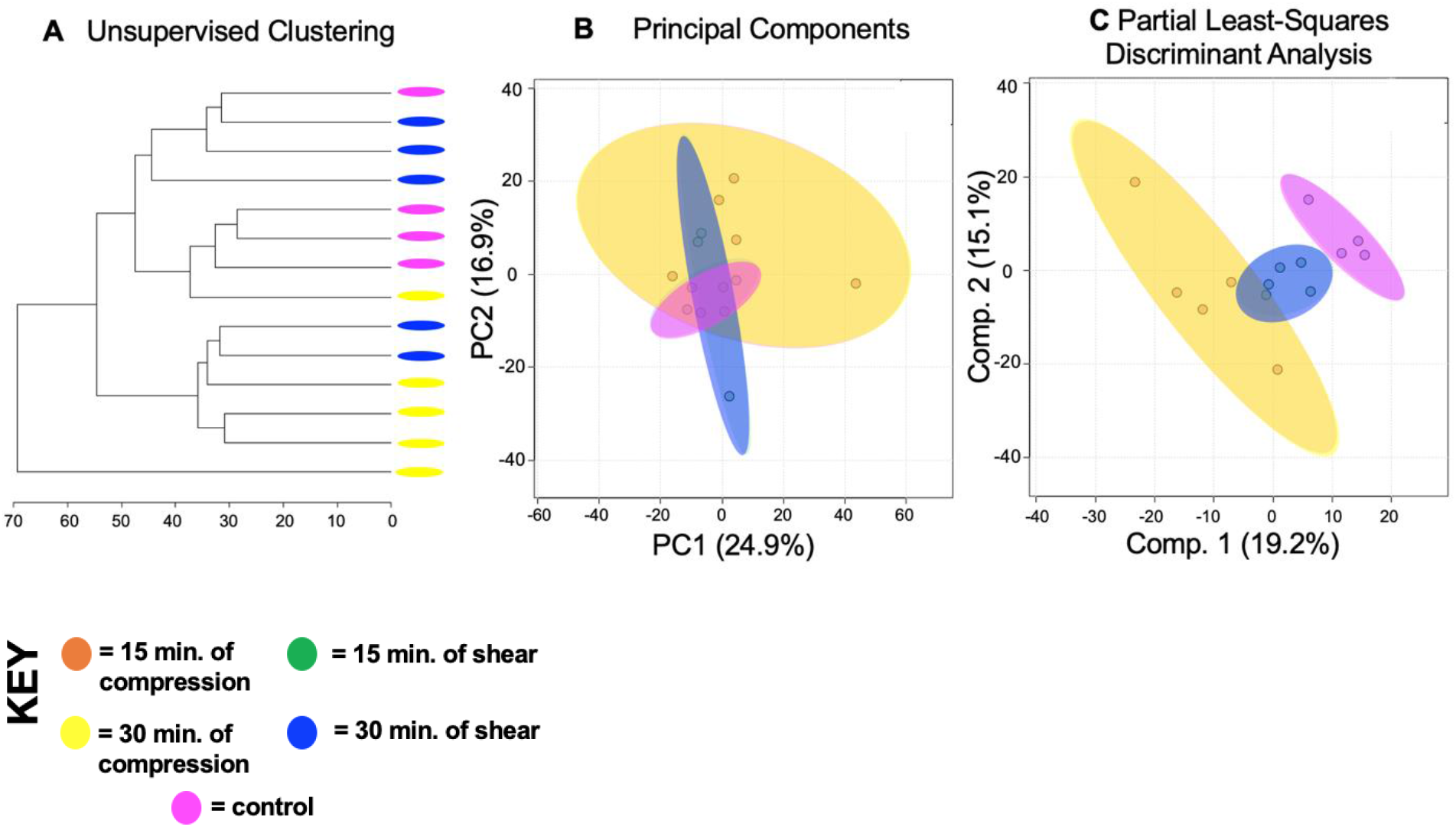
Metabolic profiles of differing forces with an equal exposure time of 30 minutes vary. Metabolomic profiles of chondrocytes exposed to compression and shear forces for 30 minutes display distinct metabolomic profiles (A). Samples exposed to different forces for the same amount of time somewhat separate into clusters by HCA as illustrated in the dendrogram. PCA displays clear clustering of samples within their respective cohorts: chondrocytes exposed to compression for 30 minutes (yellow) and chondrocytes exposed to shear forces for 30 minutes (blue). Control chondrocytes that weren’t exposed to mechanical stimuli is displayed for comparison purposes (pink). PCA is shown as a scatterplot of the first two PCs (PC1 and PC2), which account for 24.9% and 16.9% of the overall variation in the dataset respectively. PLS-DA finds clear separation between samples. PLS-DA is visualized as a scatterplot of the top two components, which account for 19.2% and 15.1% of the overall variation in the dataset.

Co-regulated metabolite features and associated pathways between shear and compressive stimulation were identified using heatmap analysis. Using the median metabolite intensities and HCA, 7 clusters were identified based on Euclidian distance. Metabolites from these clusters were then used to determine associated pathways (Fig. 6). In total, 372 pathways were detected, and 80 of the 372 are statistically significant (p < 0.05).

**Figure 6.**
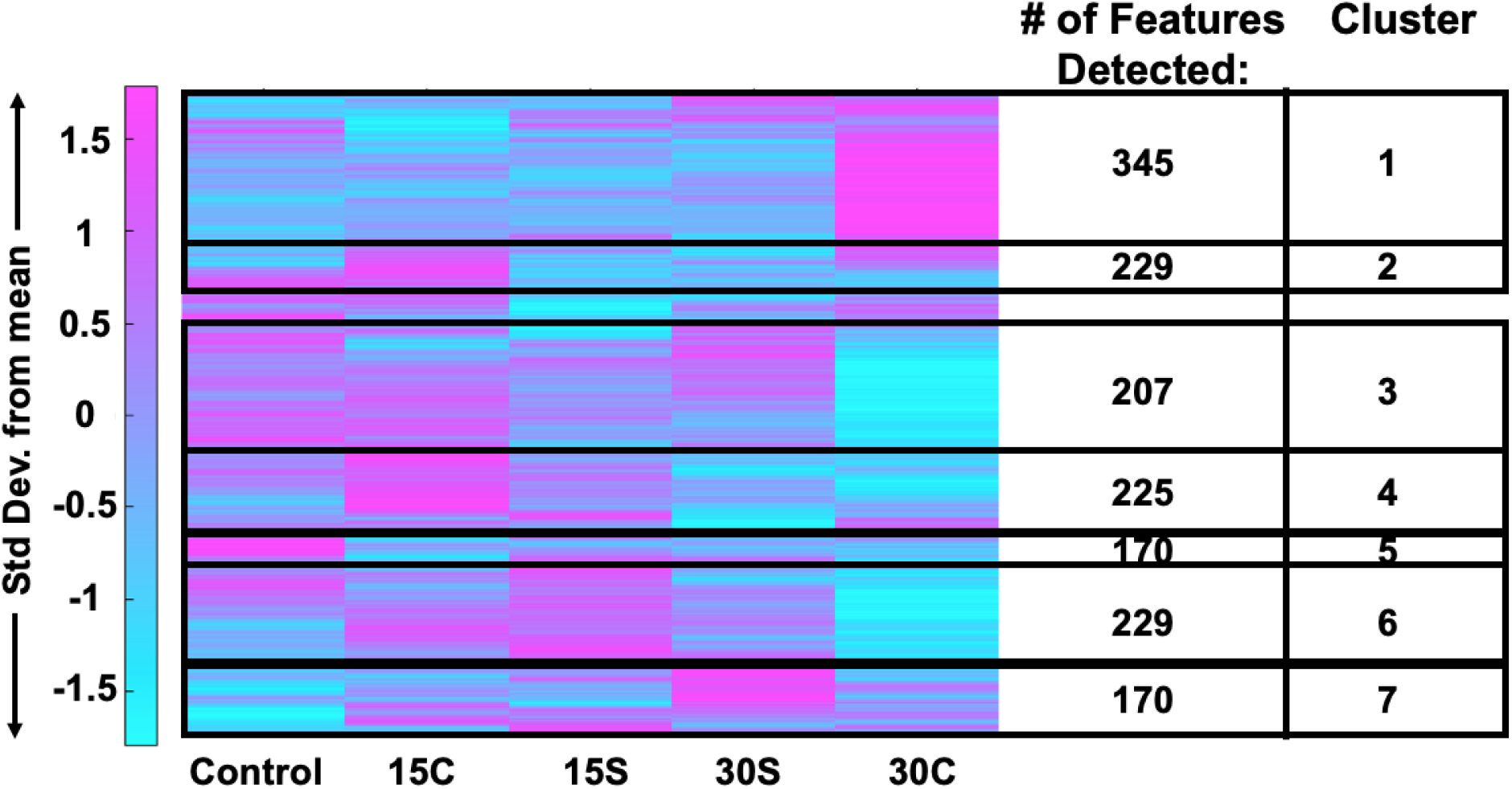
Heatmap analysis reveals that metabolic phenotypes differ when exposure time and mechanical force differ. Median intensities were clustered into 5 groups: control samples (n=4), 15 minutes of compression (n=5), 30 minutes of compression (n=5), 15 minutes of shear (n=5) and 30 minutes of shear force (n=5). 1,457 metabolite intensities were detected in SW1353 cells. To determine associated pathways and metabolites, *MS Peaks to Pathways* was employed.

Cluster 1 contained 354 metabolite features that were upregulated in samples exposed to 30 minutes of compression. These mapped to 25 statistically significant enriched pathways including primary bile acid biosynthesis, histidine metabolism, fructose and mannose metabolism, pyrimidine and purine metabolism, and alanine, aspartate, and glutamine metabolism. Cluster 2 and 4 contained 229 and 225 metabolite features, respectively, that were upregulated in samples exposed to 15 minutes of compression. These features corresponded to 23 statistically significant enriched pathways including various amino acid metabolic pathways.

Cluster 3 contained 207 metabolite features that were downregulated in samples exposed to 30 minutes of compression. These mapped to 7 statistically significant enriched pathways including alanine, aspartate, and glutamate metabolism, N-glycan biosynthesis, mannose type O-glycan biosynthesis, arginine biosynthesis, steroid hormone biosynthesis, and pyrimidine metabolism.

Cluster 5 containing 170 metabolite features that were upregulated in control samples. These features corresponded to 8 statistically significant enriched pathways including butanoate metabolism, sphingolipid metabolism, propanoate metabolism, the citrate cycle (TCA cycle), histidine metabolism, and various amino acid metabolic pathways.

Clusters 6 and 7 contained 229 and 170 metabolites that corresponded to samples exposed to shear for both 15 and 30 minutes. The features of the two combined clusters corresponded to 17 statistically significant enriched pathways including various amino acid metabolic pathways, the pentose phosphate pathway, glycolysis and gluconeogenesis.

Amino acid metabolic pathways unique to compressed SW1353 cells include alanine, asparagine, aspartate, glutamate, glutamine, histidine, isoleucine, leucine, and valine metabolism. Metabolic pathways shared by both SW1353 cells exposed to compression and shear included arginine, cysteine, glycine, lysine, methionine, proline, serine and threonine. Amino acids that weren’t detected within this dataset include phenylalanine, tryptophan, and tyrosine.

## Discussion

Cartilage provides a smooth surface for movement, reduces stress placed on underlying bone, and requires a strong and smooth ECM to resist high loads created in the joint. To further investigate the role of cartilage and mechanotransduction in chondrocytes, we investigated the effects of mechanical stimuli on the ECM and how differing types of load (shear and compression) and exposure times affect the metabolism of SW1353 chondrocytes. In total, 1,457 metabolites were detected, and metabolic profiles were generated for each experimental condition (time pt. 0, 15, 30 min., mechanical stimulus - shear or compression).

In every pairwise comparison, metabolomic profiles from different experimental groups differed from each other. When comparing all experimental groups, sialic acid metabolism was the most notable pathway determined from the top 100 VIP metabolites when comparing all groups. This finding corresponds to sialic acid being an inflammatory biomarker for OA and rheumatoid arthritis (RA), while also contributing to the lubrication of the joint (Alturfan et al., 2007). Additionally, lipid metabolism (fatty acid metabolism, cholesterol biosynthesis, prostaglandin formation from arachidonate) was upregulated in compressed samples. Based on potential crosstalk between joint tissues, the permeability of AC, and the ability of fats like phospholipids to act as lubricants, overexpression of fat associated pathways may lead to cartilage lesions, joint space narrowing, joint immobilization, and ultimately contributing to the development of OA (Appleton et al., 2007; Lotke and Granda, 1972; Maroudas et al., 1968; Pan et al., 2012; Sohn et al., 2012; Villalvilla et al., 2013). Central energy metabolism and beta-oxidation were present across experimental groups and strongly upregulated amongst compressed samples (TCA cycle, carnitine shuttle). We interpret this as exposure to mechanical stimuli driving a high chondrocyte ATP demand: the TCA cycle and the carnitine shuttle may have been upregulated to meet this need. Amino acid metabolism differed between samples exposed to shear and compression while several specific amino acids were unique to compression. Taken together, compression may induce mechanosensitive pathways that are needed to produce a broader set of products than shear such as amino acids, both non-essential and essential.

Many studies have shown the diverse effects of mechanical stimulation on the musculoskeletal system, chondrocytes, ECM, cell metabolism, and its relation to OA (Jutila et al., 2014; Jutila et al., 2015; Salinas et al., 2017; Zignego et al., 2015). Bushmann *et al* investigated the effects of static and dynamic mechanical compression on matrix biosynthesis, chondrocyte proliferation, and quantified proteoglycan and glycosaminoglycan content. Through the use of cell culture and radiolabeling, *in vitro* chondrocytes form a mechanically functional matrix that preserves certain physiological features of chondrocyte behavior and response to physical stimuli (Buschmann et al., 1995; Buschmann et al., 1992). O’Connor *et al* examined the effects of mechanical stimulation on chondrocyte metabolism and transient receptor potential vanilloid 4 (TRPV4). The results of this study suggest that dynamic loading has a profound effect on cell physiology, ion channel function, and TRPV4 mediated mechanotransduction (O’Conor et al., 2014).

Jutila *et al* report the effects of compressive loading at early time points (0, 15 and 30 minutes). Through the utilization of targeted and untargeted metabolomics, 54 metabolites were found to mediate chondrocyte mechanotransduction (Jutila et al., 2014; Jutila et al., 2015). Other studies identified the effects of physiological compression on energy metabolism and maintenance of the pericellular and extracellular matrices (Salinas et al., 2017; Zignego et al., 2015). In addition, many pathways related to the metabolism of energy, lipids, and amino acids were identified through targeted and untargeted metabolomics. The results of our study are consistent with these prior studies and together indicate that extended exposure to mechanical stimuli affects chondrocytes, ECM components, and alters cellular metabolism.

### Inflammation

Exposure to shear and compression lead to metabolic shifts associated with inflammation and deterioration in SW1353 cells. The 100 most significant metabolites across all sample groups were involved in sialic acid metabolism (Fig. 1). Sialic acid (SA) is an acylated derivative of neuraminic acid attached to glycoproteins and glycolipids. Serum SA levels are a known marker of inflammation and has been reported as a useful biomarker of inflammation in patients with OA and RA (Cui et al., 2014). Additionally, studies find that differing levels of SA are linked to OA severity and could be used as a diagnostic tool in the future (Alturfan et al., 2007; Browning et al., 2004; Chavan et al., 2005; Cui et al., 2014; Gobezie et al., 2007; Kosakai, 1991). Our study provides further evidence that sialic acid production could be a potential marker for OA due to its metabolic presence when mechanically stimulated with shear and compressive forces. Sialic acid is also a key component of the superficial zone protein lubricin that decreases cartilage-on-cartilage friction (Carlson et al., 2019; Jay et al., 2012; Jutila et al., 2014; Jutila et al., 2015). Therefore, these findings may suggest different loading types and exposure times may promote changes in metabolism that correspond to changes in cartilage structure and function. Taken together, this is the first study to find sialic acid metabolism to be upregulated after mechanical stimuli and further research may identify its effects on the ECM, cartilage health, and OA.

Further analysis showed that prostaglandin formation from arachidonate was a significant pathway associated with compression. Prostaglandin metabolites were upregulated after 30 minutes of compression. Studies show that prostaglandin formation from arachidonate can be triggered by obesity, age, and mechanical stress (Bar-Or et al., 2015). Individually or combined, these stimuli result in cartilage deterioration which can be attributed to the innate immune system, by pathogen-associated molecular patterns (PAMPs), damage-associated molecular patterns (DAMPs), and alarmins (Bianchi, 2007). Alarmins are triggered by signals from cellular damage, abnormal proteins, leaky vasculature, and fragments of cartilage matrix (Bar-Or et al., 2015; Bianchi, 2007). Prostaglandins such as PGD2, PGE2, and others are anti-inflammatory molecules that initiate a decrease in inflammation and trigger recovery to normal cellular function (Attur et al., 2008; Wang et al., 2013; Wang et al., 2010). Upon the application of low fluid shear stress anti-inflammatory molecules are activated, inflammation is halted, and recovery is initiated (Attur et al., 2008; Bar-Or et al., 2015; Wang et al., 2013). The results of this study suggest that increased compressive loading leads to an increased presence of inflammatory pathways such as prostaglandin formation from arachidonate in SW1353 cells. Therefore, if confirmed in primary cells, increased compressive stimulation may result in increased inflammation, which in excess could lead to OA if not balanced by other physiological processes.

### Lipid Metabolism

In this study, a common metabolic theme of SW1353 cells stimulated by compression was upregulation of metabolite features corresponding to lipid metabolism. The pathways detected include fatty acid biosynthesis, cholesterol biosynthesis, and prostaglandin formation from arachidonate which were significantly upregulated after 30 minutes of compression. Further, butanoate and sphingolipid metabolism were significantly upregulated in control samples, implying downregulation upon mechanical stimulation.

When joint homeostasis is altered, increased protease activity can result in cartilage lesions, joint space narrowing, and ultimately, breakdown of the tissues (Appleton et al., 2007; Lotke and Granda, 1972; Maroudas et al., 1968; Pan et al., 2012; Sohn et al., 2012; Villalvilla et al., 2013). Hence cartilage health and metabolism greatly rely on the chondrocyte microenvironment. On the metabolic level, both fatty acid metabolism and cholesterol biosynthesis are required to maintain healthy ECM and cartilage (Aguilar et al., 2009; Bernstein et al., 2010; Gkretsi et al., 2011; Wu and De Luca, 2004). Cartilage contains stores of lipid deposits, especially in chondrocytes, during times of health and disease. Expected lipid content in both healthy and pathological cartilage include palmitic, linolic, and oleic acid, but fatty acids and arachidonate acid are elevated in OA samples and associated with increased histological severity (Lippiello et al., 1991). Beyond prostaglandin synthesis from arachidonate, in this study both fatty acid metabolism and cholesterol biosynthesis were upregulated in loaded samples indicating that chondrocyte metabolism was altered as a result of shear and compression stimulation. Future studies may determine the role of mechanical stimuli in regulating the cartilage ECM through these pathways.

### Central Metabolism

Injurious mechanical stimuli, traumatic injury, and others factors such as obesity, age, and gender initiate the breakdown of cartilage leading to OA. In contrast, lower levels of mechanical stimuli result in matrix synthesis to support cartilage homeostasis. Both repair and maintenance require additional energy production to produce ATP. Pathways that generate ATP are pathways related to central metabolism including glycolysis, the TCA cycle, beta-oxidation, and the carnitine shuttle. In this study, pathways related to the glycolysis, TCA cycle, and the carnitine shuttle were significantly upregulated consistent with prior results (Salinas et al., 2017; Salinas et al., 2019; Zignego et al., 2015).

The carnitine shuttle was upregulated in all mechanically stimulated samples after 15 minutes of either compression or shear. Beta-oxidation breaks down fatty acids to produce energy and is regulated by the carnitine shuttle. Carnitines stimulate cell proliferation, induce ATP synthesis, and serum carnitines were associated with OA grade (Tootsi et al., 2018; Zhai, 2019).

Metabolites and corresponding pathways involved in glycolysis and the TCA cycle were upregulated in SW1353 cells exposed to 15 minutes of shear. In chondrocytes, anaerobic glycolysis is restricted as oxygen and nutrients are limited in avascular cartilage. But previous studies have found that both aerobic and anaerobic glycolysis occur in chondrocytes (Zheng et al., 2021). Specific to OA, the rate of anerobic glycolysis in diseased cartilage is higher compared to healthy cartilage (Maneiro et al., 2003). Other intermediates that can be incorporated into glycolysis to yield energy are fructose, mannose, and galactose. In this study, fructose, mannose, and galactose metabolism were upregulated in SW1353 cells exposed to compression.

The TCA cycle produces ATP from pyruvate and other sources such as sugar, fat, and proteins. Intermediate TCA metabolites include citrate, malate, succinate, fumarate, and various others. Studies find that patients with OA have higher levels of synovium metabolites associated with the TCA cycle compared to healthy controls (Adams et al., 2012; Anderson et al., 2018; Zhai, 2019). The results of this study provide evidence that mechanical stimulation by compression specifically generates additional ATP. This ATP may be used to maintain and produce matrix thus requiring upregulation of the TCA cycle and the carnitine shuttle.

Beyond glycolysis and the TCA cycle, the pentose phosphate pathway (PPP) was upregulated in SW1353 cells exposed to compression. The PPP pathway utilizes glucose-6-phosphate which is a product of glycolysis. The PPP gives rise to NADH and ribose-5-phosphate which is used to make subunits for DNA, RNA, and amino acids such as histidine from phosphoribosyl pyrophosphate (PRPP). Here, histidine and beta-alanine metabolism were upregulated only in SW1353 cells exposed to compression. Histamine is the decarboxylated amine form of histidine, and previous studies have found that histamines stimulate chondrocytes proliferation in humans (Tetlow and Woolley, 2003; Tetlow and Woolley, 2005). Therefore, the upregulation of histidine metabolism in this study suggests that SW1353 cells exposed to compression may proliferate in response mechanical stimuli.

Finally, two pathways detected in SW1353 cells stimulated by 30 minutes of compression were pyrimidine and purine metabolism (p-value < 0.05). Pyrimidine and purine metabolism stem from the pentose phosphate pathway. These two pathways function to recycle nucleosides and participate in *de novo* nucleotide synthesis. Upregulation of these pathways may relate to an increase in cellular demand of complex molecules such as RNA upon mechanical stimulation (e.g. to support transcription). Beyond nucleotide and nucleoside production, pyrimidine and purine metabolism give rise to various amino acids. In particular, pyrimidine interconversions can lead to beta-alanine metabolism. Previous studies have found increased levels of beta-alanine in urine, synovium, synovial fluid, and in subchondral bone in OA animal models (Hugle et al., 2012; Maher et al., 2012; Okun et al., 2002; Yang et al., 2016).

### Amino Acid Metabolism

Metabolomic profiles for SW1353 cells stimulated by either shear or compression show that amino acid metabolism is commonly upregulated by these types of loading with certain responses specific to the type and/or duration of loading. Amino acid pathways unique to compressive stimulation include metabolism of alanine, asparagine, aspartate, glutamate, glutamine, histidine, isoleucine, leucine, and valine metabolism. In contrast, arginine, cysteine, glycine, lysine, methionine, proline, serine, and threonine were induced by both shear and stimuli compression (Fig. 7).

**Figure 7.**
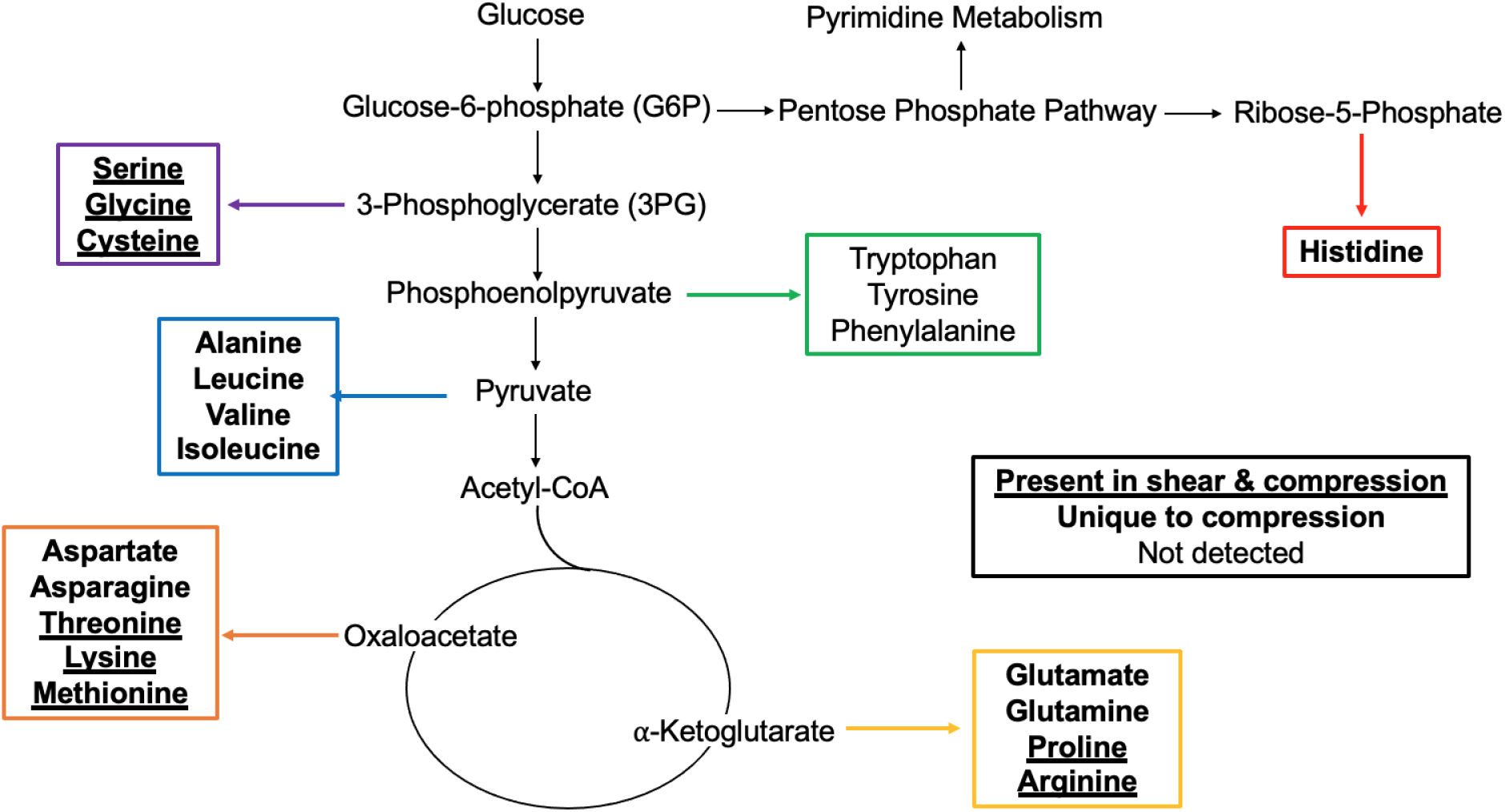
Mapping of mechanically stimulated amino acids in metabolic pathways. Amino acids that are bolded and underlined were upregulated in SW1353 cells that were exposed to compression and shear force. Those bolded only were upregulated in cells exposed to compression. Amino acids that aren’t bolded nor underlined weren’t detected (tryptophan, tyrosine, phenylalanine). Intermediates of interest include ribose-5-phosphate, 3-phosphoglycerate, pyruvate, α-ketoglutarate and oxaloacetate. Additionally, pyrimidine metabolism was upregulated in samples exposed to 30 minutes of compression which stems from the pentose phosphate pathway.

Amino acids are the building blocks of proteins and determine structure and function. Depending on cellular demand, amino acids can be converted into intermediates, or produced from glucose, through central energy metabolism. The results of this study suggest that major precursors of interest when analyzing the relationship between mechanical stimuli and metabolism include ribose-5-phosphate (histidine), 3-phosphoglycerate (serine, glycine, cysteine), pyruvate (alanine, leucine, valine, isoleucine), α-ketoglutarate (glutamate, glutamine, proline, arginine) and oxaloacetate (aspartate, asparagine, threonine, lysine, methionine) (Fig. 7). The results of this study suggest that the upregulation of metabolic pathways may correspond to the synthesis of essential and non-essential amino acids that are needed to synthesize musculoskeletal associated proteins in response to mechanical stimuli. Furthermore, the application of different stimuli, such as compression and shear, results in different metabolic profiles that impact protein synthesis of various proteins.

### Limitations

This study has important limitations and opportunities for future studies. First, the behavior of the chondrocyte cell line SW1353 cells may differ from primary human chondrocytes. Second, although our *in vitro* approach is required to apply well-defined mechanical loads, this approach may yield different results from an *in vivo* model or one where the pericellular matrix is present. Third, exposure to mechanical stimuli for both periods shorter than 15 minutes and beyond 30 minutes could expand our knowledge on the varying metabolic phenotypes beyond the those in this study. Finally, this was an untargeted metabolomic study. By performing a more targeted metabolomic analysis utilizing MS/MS, uncertainty of predicted identities of key metabolite features and their associated pathways can be reduced.

## Conclusion

To our knowledge, this is the first study to generate global metabolomic profiles of SW1353 cells exposed to both shear and compression at early time points. Pathways were identified within experimental groups which provide insight into the role of the extracellular matrix, articular cartilage homeostasis, and the potential outcomes of metabolic alterations that ultimately lead to osteoarthritis. Furthermore, these metabolomic profiles support the link between mechanical stimuli and cartilage remodeling. Expansion of this study may identify cartilage loading protocols to bolster matrix synthesis that may be relevant to drug development.

**Table 1.** Amino acid metabolic pathways detected in SW1353 cells exposed to mechanical stimuli. Untargeted metabolomic analyses reveal that amino acid metabolism is a common metabolic theme in samples exposed to shear and compressive forces. Amino acids unique to SW1353 cells exposed to compression include alanine, asparagine, aspartate, glutamate, glutamine, histidine, isoleucine, leucine, and valine.

| Amino Acid | Mechanical Stimuli |  |
| --- | --- | --- |
|  | Compression | Shear |
| Alanine | X |  |
| Arginine | X | X |
| Asparagine | X |  |
| Aspartate | X |  |
| Cysteine | X | X |
| Glutamate | X |  |
| Glutamine | X |  |
| Glycine | X | X |
| Histidine | X |  |
| Isoleucine | X |  |
| Leucine | X |  |
| Lysine | X | X |
| Methionine | X | X |
| Phenylalanine |  |  |
| Proline | X | X |
| Serine | X | X |
| Threonine | X | X |
| Tryptophan |  |  |
| Tyrosine |  |  |
| Valine | X |  |

## Abbreviations List

ECM: extracellular matrix
OA: osteoarthritis
LC-MS: liquid chromatography-mass spectrometry
HPLC-MS: high precision liquid chromatography and mass spectrometry
HCA: hierarchical cluster analysis
PCA: principal component analysis
PC: principal components
PLS-DA: partial least-squares discriminant analysis
VIP: variable important in projection
FDR: false discovery rate
M/Z: mass-to-charge ratio

## Acknowledgements

The authors gratefully acknowledge funding from NSF (CMMI 1554708) and NIH (R01AR073964). Funding for the Proteomics, Metabolomics and Mass Spectrometry Facility was made possible in part by the MJ Murdock Charitable Trust and the National Institute of General Medical Sciences of the National Institutes of Health under Award Number P20GM103474. The content is solely the responsibility of the authors and does not necessarily represent the official views of the National Institutes of Health.

## Author contributions

CNM performed cell culturing and mechanical stimulation of SW1353 cells. HDW analyzed data and drafted the manuscript. RKJ designed experiments and assisted in analyzing data. All authors have read and revised the manuscript.

## Supplemental Materials and Methods

### Statistical Analysis for Untargeted Metabolomic Profiling

Univariate, supervised, and unsupervised multivariate analyses were used to visualize and narrow the dataset. The unsupervised multivariate statistical analyses that were utilized were hierarchical cluster analysis (HCA) and principal component analysis (PCA). HCA is used to visualize metabolomic profiles, identify sub-groups of sample sets, and determine differences between experimental groups. PCA linearly transforms and reduces the high dimensionality dataset into latent variables – specifically principal components (PCs) – to explain the variability in the dataset. Each PC is a combination of metabolite features that contributed most to the clustering of samples.

To further visualize the dataset and identify specific differences between cohorts, we used partial least-squares discriminant analysis (PLS-DA), variable importance in projection scores (VIP) and volcano plot analysis. PLS-DA is similar to PCA but is a supervised statistical analysis that is utilized to seek out differences between cohorts and reveal metabolites that are contributing to the separation between cohorts. PLS-DA is especially useful to this study because it assigns variable importance in projection (VIP) scores to metabolites that are the most important in discriminating between cohorts. Volcano plot analysis was also utilized to identify significantly upregulated and downregulated metabolites with a significance level of 0.05 and false discovery rate (FDR) corrections were applied to correct for multiple comparisons. Specific metabolite features of interest identified by VIP scores and volcano plot analysis were then matched to metabolite identities and metabolic pathways using MetaboAnalyst.

**Supplementary Figure 1.**
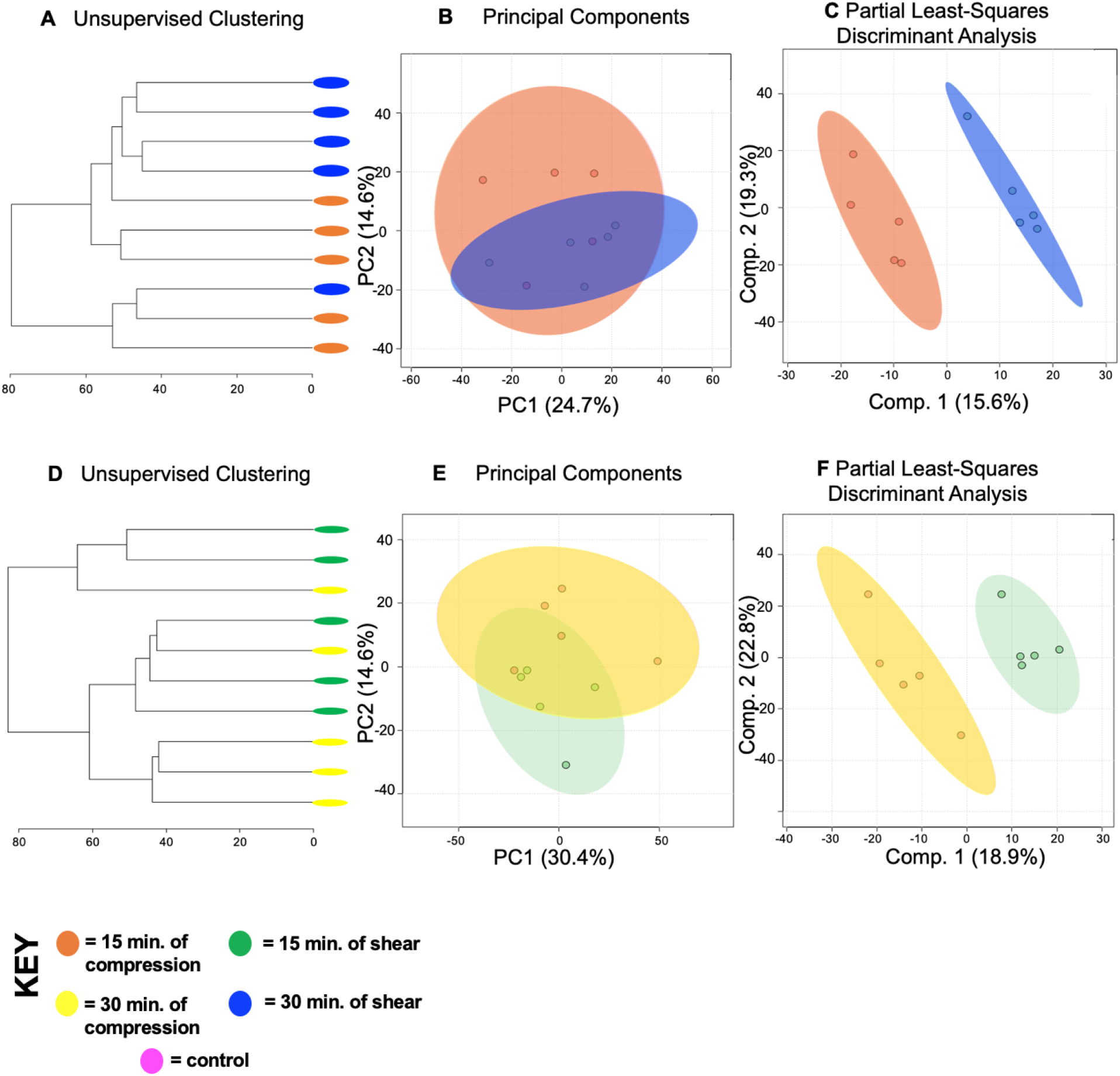
Metabolic profiles of differing forces with different exposure times vary. Metabolomic profiles of chondrocytes exposed to 15 minutes of compression and 30 minutes of shear forces display distinct metabolomic profiles (A). Samples exposed to different forces for differing amounts of time somewhat separate into clusters by HCA as illustrated in the dendrogram. (B) PCA displays clear clustering of samples within their respective cohorts: chondrocytes exposed to compression for 15 minutes (orange) and chondrocytes exposed to shear forces for 30 minutes (blue). PCA is shown as a scatterplot of the first two PCs (PC1 and PC2), which account for 24.7% and 14.6% of the overall variation in the dataset respectively. (C) PLS-DA finds clear separation between samples. PLS-DA is visualized as a scatterplot of the top two components, which account for 15.6% and 19.3% of the overall variation in the dataset. Additionally, metabolomic profiles of chondrocytes exposed to 30 minutes of compression and 15 minutes of shear force are distinct from each other. (D) HCA illustrates that groups somewhat separate. (E) PCA displays overlap of samples between chondrocytes exposed to different forces and exposure times. PC1 and PC2 account for 30.4% and 14.6% of the overall variation in the dataset respectively. (F) PLS-DA finds clear separation between samples. The two components account for 18.9% and 22.8% of variation in the dataset.

## References

Adams, S. B., Jr., Setton, L. A., Kensicki, E., Bolognesi, M. P., Toth, A. P. and Nettles, D. L. (2012). Global metabolic profiling of human osteoarthritic synovium. Osteoarthritis Cartilage 20, 64–7.

Aguilar, A., Wu, S. and De Luca, F. (2009). P450 oxidoreductase expressed in rat chondrocytes modulates chondrogenesis via cholesterol- and Indian Hedgehog-dependent mechanisms. Endocrinology 150, 2732–9.

Alturfan, A. A., Uslu, E., Alturfan, E. E., Hatemi, G., Fresko, I. and Kokoglu, E. (2007). Increased serum sialic acid levels in primary osteoarthritis and inactive rheumatoid arthritis. Tohoku J Exp Med 213, 241–8.

Anderson, J. R., Chokesuwattanaskul, S., Phelan, M. M., Welting, T. J. M., Lian, L. Y., Peffers, M. J. and Wright, H. L. (2018). (1)H NMR Metabolomics Identifies Underlying Inflammatory Pathology in Osteoarthritis and Rheumatoid Arthritis Synovial Joints. J Proteome Res 17, 3780–3790.

Appleton, C. T., Pitelka, V., Henry, J. and Beier, F. (2007). Global analyses of gene expression in early experimental osteoarthritis. Arthritis Rheum 56, 1854–68.

Attur, M., Al-Mussawir, H. E., Patel, J., Kitay, A., Dave, M., Palmer, G., Pillinger, M. H. and Abramson, S. B. (2008). Prostaglandin E2 exerts catabolic effects in osteoarthritis cartilage: evidence for signaling via the EP4 receptor. J Immunol 181, 5082–8.

Bar-Or, D., Rael, L. T., Thomas, G. W. and Brody, E. N. (2015). Inflammatory Pathways in Knee Osteoarthritis: Potential Targets for Treatment. Curr Rheumatol Rev 11, 50–58.

Bernstein, P., Sticht, C., Jacobi, A., Liebers, C., Manthey, S. and Stiehler, M. (2010). Expression pattern differences between osteoarthritic chondrocytes and mesenchymal stem cells during chondrogenic differentiation. Osteoarthritis Cartilage 18, 1596–607.

Bianchi, M. E. (2007). DAMPs, PAMPs and alarmins: all we need to know about danger. J Leukoc Biol 81, 1–5.

Bricca, A., Juhl, C. B., Grodzinsky, A. J. and Roos, E. M. (2017). Impact of a daily exercise dose on knee joint cartilage - a systematic review and meta-analysis of randomized controlled trials in healthy animals. Osteoarthritis Cartilage 25, 1223–1237.

Browning, L. M., Krebs, J. D. and Jebb, S. A. (2004). Discrimination ratio analysis of inflammatory markers: implications for the study of inflammation in chronic disease. Metabolism 53, 899–903.

Buschmann, M. D., Gluzband, Y. A., Grodzinsky, A. J. and Hunziker, E. B. (1995). Mechanical compression modulates matrix biosynthesis in chondrocyte/agarose culture. J Cell Sci 108 (Pt 4), 1497–508.

Buschmann, M. D., Gluzband, Y. A., Grodzinsky, A. J., Kimura, J. H. and Hunziker, E. B. (1992). Chondrocytes in agarose culture synthesize a mechanically functional extracellular matrix. J Orthop Res 10, 745–58.

Carlson, A. K., Rawle, R. A., Adams, E., Greenwood, M. C., Bothner, B. and June, R. K. (2018). Application of global metabolomic profiling of synovial fluid for osteoarthritis biomarkers. Biochem Biophys Res Commun 499, 182–188.

Carlson, A. K., Rawle, R. A., Wallace, C. W., Brooks, E. G., Adams, E., Greenwood, M. C., Olmer, M., Lotz, M. K., Bothner, B. and June, R. K. (2019). Characterization of synovial fluid metabolomic phenotypes of cartilage morphological changes associated with osteoarthritis. Osteoarthritis Cartilage 27, 1174–1184.

Chan, D. D., Cai, L., Butz, K. D., Nauman, E. A., Dickerson, D. A., Jonkers, I. and Neu, C. P. (2018). Functional MRI can detect changes in intratissue strains in a full thickness and critical sized ovine cartilage defect model. J Biomech 66, 18–25.

Chan, D. D., Cai, L., Butz, K. D., Trippel, S. B., Nauman, E. A. and Neu, C. P. (2016). In vivo articular cartilage deformation: noninvasive quantification of intratissue strain during joint contact in the human knee. Sci Rep 6, 19220.

Chavan, M. M., Kawle, P. D. and Mehta, N. G. (2005). Increased sialylation and defucosylation of plasma proteins are early events in the acute phase response. Glycobiology 15, 838–48.

Cui, Z., Liu, K., Wang, A., Liu, S., Wang, F. and Li, J. (2014). Correlation between sialic acid levels in the synovial fluid and the radiographic severity of knee osteoarthritis. Exp Ther Med 8, 255–259.

Gillespie, P. G. and Muller, U. (2009). Mechanotransduction by hair cells: models, molecules, and mechanisms. Cell 139, 33–44.

Gkretsi, V., Simopoulou, T. and Tsezou, A. (2011). Lipid metabolism and osteoarthritis: lessons from atherosclerosis. Prog Lipid Res 50, 133–40.

Gobezie, R., Kho, A., Krastins, B., Sarracino, D. A., Thornhill, T. S., Chase, M., Millett, P. J. and Lee, D. M. (2007). High abundance synovial fluid proteome: distinct profiles in health and osteoarthritis. Arthritis Res Ther 9, R36.

Hugle, T., Kovacs, H., Heijnen, I. A., Daikeler, T., Baisch, U., Hicks, J. M. and Valderrabano, V. (2012). Synovial fluid metabolomics in different forms of arthritis assessed by nuclear magnetic resonance spectroscopy. Clin Exp Rheumatol 30, 240–5.

Jaalouk, D. E. and Lammerding, J. (2009). Mechanotransduction gone awry. Nat Rev Mol Cell Biol 10, 63–73.

Jay, G. D., Elsaid, K. A., Kelly, K. A., Anderson, S. C., Zhang, L., Teeple, E., Waller, K. and Fleming, B. C. (2012). Prevention of cartilage degeneration and gait asymmetry by lubricin tribosupplementation in the rat following anterior cruciate ligament transection. Arthritis Rheum 64, 1162–71.

Jewett, M. C., Hofmann, G. and Nielsen, J. (2006). Fungal metabolite analysis in genomics and phenomics. Curr Opin Biotechnol 17, 191–7.

Jutila, A. A., Zignego, D. L., Hwang, B. K., Hilmer, J. K., Hamerly, T., Minor, C. A., Walk, S. T. and June, R. K. (2014). Candidate mediators of chondrocyte mechanotransduction via targeted and untargeted metabolomic measurements. Arch Biochem Biophys 545, 116–23.

Jutila, A. A., Zignego, D. L., Schell, W. J. and June, R. K. (2015). Encapsulation of chondrocytes in high-stiffness agarose microenvironments for in vitro modeling of osteoarthritis mechanotransduction. Ann Biomed Eng 43, 1132–44.

Kosakai, O. (1991). [Clinical relevance of sialic acids determination in serum and synovial fluid in orthopaedic disorders]. Rinsho Byori 39, 197–207.

Lippiello, L., Walsh, T. and Fienhold, M. (1991). The association of lipid abnormalities with tissue pathology in human osteoarthritic articular cartilage. Metabolism 40, 571–6.

Lotke, P. A. and Granda, J. L. (1972). Alterations in the permeability of articular cartilage by proteolytic enzymes. Arthritis Rheum 15, 302–8.

Maher, A. D., Coles, C., White, J., Bateman, J. F., Fuller, E. S., Burkhardt, D., Little, C. B., Cake, M., Read, R., McDonagh, M. B. et al. (2012). 1H NMR spectroscopy of serum reveals unique metabolic fingerprints associated with subtypes of surgically induced osteoarthritis in sheep. J Proteome Res 11, 4261–8.

Maneiro, E., Martin, M. A., de Andres, M. C., Lopez-Armada, M. J., Fernandez-Sueiro, J. L., del Hoyo, P., Galdo, F., Arenas, J. and Blanco, F. J. (2003). Mitochondrial respiratory activity is altered in osteoarthritic human articular chondrocytes. Arthritis Rheum 48, 700–8.

Maroudas, A., Bullough, P., Swanson, S. A. and Freeman, M. A. (1968). The permeability of articular cartilage. J Bone Joint Surg Br 50, 166–77.

O’Conor, C. J., Leddy, H. A., Benefield, H. C., Liedtke, W. B. and Guilak, F. (2014). TRPV4-mediated mechanotransduction regulates the metabolic response of chondrocytes to dynamic loading. Proc Natl Acad Sci U S A 111, 1316–21.

Okun, J. G., Kolker, S., Schulze, A., Kohlmuller, D., Olgemoller, K., Lindner, M., Hoffmann, G. F., Wanders, R. J. and Mayatepek, E. (2002). A method for quantitative acylcarnitine profiling in human skin fibroblasts using unlabelled palmitic acid: diagnosis of fatty acid oxidation disorders and differentiation between biochemical phenotypes of MCAD deficiency. Biochim Biophys Acta 1584, 91–8.

Orr, A. W., Helmke, B. P., Blackman, B. R. and Schwartz, M. A. (2006). Mechanisms of mechanotransduction. Dev Cell 10, 11–20.

Paluch, E. K., Nelson, C. M., Biais, N., Fabry, B., Moeller, J., Pruitt, B. L., Wollnik, C., Kudryasheva, G., Rehfeldt, F. and Federle, W. (2015). Mechanotransduction: use the force(s). BMC Biol 13, 47.

Pan, J., Wang, B., Li, W., Zhou, X., Scherr, T., Yang, Y., Price, C. and Wang, L. (2012). Elevated cross-talk between subchondral bone and cartilage in osteoarthritic joints. Bone 51, 212–7.

Racunica, T. L., Teichtahl, A. J., Wang, Y., Wluka, A. E., English, D. R., Giles, G. G., O’Sullivan, R. and Cicuttini, F. M. (2007). Effect of physical activity on articular knee joint structures in community-based adults. Arthritis Rheum 57, 1261–8.

Salinas, D., Minor, C. A., Carlson, R. P., McCutchen, C. N., Mumey, B. M. and June, R. K. (2017). Combining Targeted Metabolomic Data with a Model of Glucose Metabolism: Toward Progress in Chondrocyte Mechanotransduction. PLoS One 12, e0168326.

Salinas, D., Mumey, B. M. and June, R. K. (2019). Physiological dynamic compression regulates central energy metabolism in primary human chondrocytes. Biomech Model Mechanobiol 18, 69–77.

Sanchez-Adams, J., Leddy, H. A., McNulty, A. L., O’Conor, C. J. and Guilak, F. (2014). The mechanobiology of articular cartilage: bearing the burden of osteoarthritis. Curr Rheumatol Rep 16, 451.

Sohn, D. H., Sokolove, J., Sharpe, O., Erhart, J. C., Chandra, P. E., Lahey, L. J., Lindstrom, T. M., Hwang, I., Boyer, K. A., Andriacchi, T. P. et al. (2012). Plasma proteins present in osteoarthritic synovial fluid can stimulate cytokine production via Toll-like receptor 4. Arthritis Res Ther 14, R7.

Sophia Fox, A. J., Bedi, A. and Rodeo, S. A. (2009). The basic science of articular cartilage: structure, composition, and function. Sports Health 1, 461–8.

Tetlow, L. C. and Woolley, D. E. (2003). Histamine stimulates the proliferation of human articular chondrocytes in vitro and is expressed by chondrocytes in osteoarthritic cartilage. Ann Rheum Dis 62, 991–4.

Tetlow, L. C. and Woolley, D. E. (2005). Histamine, histamine receptors (H1 and H2), and histidine decarboxylase expression by chondrocytes of osteoarthritic cartilage: an immunohistochemical study. Rheumatol Int 26, 173–8.

Tootsi, K., Kals, J., Zilmer, M., Paapstel, K., Ottas, A. and Martson, A. (2018). Medium- and long-chain acylcarnitines are associated with osteoarthritis severity and arterial stiffness in end-stage osteoarthritis patients: a case-control study. Int J Rheum Dis 21, 1211–1218.

Villalvilla, A., Gomez, R., Largo, R. and Herrero-Beaumont, G. (2013). Lipid transport and metabolism in healthy and osteoarthritic cartilage. Int J Mol Sci 14, 20793–808.

Wang, P., Guan, P. P., Guo, C., Zhu, F., Konstantopoulos, K. and Wang, Z. Y. (2013). Fluid shear stress-induced osteoarthritis: roles of cyclooxygenase-2 and its metabolic products in inducing the expression of proinflammatory cytokines and matrix metalloproteinases. FASEB J 27, 4664–77.

Wang, P., Zhu, F., Lee, N. H. and Konstantopoulos, K. (2010). Shear-induced interleukin-6 synthesis in chondrocytes: roles of E prostanoid (EP) 2 and EP3 in cAMP/protein kinase A- and PI3-K/Akt-dependent NF-kappaB activation. J Biol Chem 285, 24793–804.

Wu, S. and De Luca, F. (2004). Role of cholesterol in the regulation of growth plate chondrogenesis and longitudinal bone growth. J Biol Chem 279, 4642–7.

Xia, J. and Wishart, D. S. (2016). Using MetaboAnalyst 3.0 for Comprehensive Metabolomics Data Analysis. Curr Protoc Bioinformatics 55, 14 10 1–14 10 91.

Yang, G., Zhang, H., Chen, T., Zhu, W., Ding, S., Xu, K., Xu, Z., Guo, Y. and Zhang, J. (2016). Metabolic analysis of osteoarthritis subchondral bone based on UPLC/Q-TOF-MS. Anal Bioanal Chem 408, 4275–86.

Zhai, G. (2019). Alteration of Metabolic Pathways in Osteoarthritis. Metabolites 9.

Zheng, L., Zhang, Z., Sheng, P. and Mobasheri, A. (2021). The role of metabolism in chondrocyte dysfunction and the progression of osteoarthritis. Ageing Res Rev 66, 101249.

Zignego, D. L., Hilmer, J. K. and June, R. K. (2015). Mechanotransduction in primary human osteoarthritic chondrocytes is mediated by metabolism of energy, lipids, and amino acids. J Biomech 48, 4253–61.

